# Hormonal regulation of Semaphorin 7a in ER+ breast cancer drives therapeutic resistance

**DOI:** 10.1101/650135

**Authors:** Lyndsey S Crump, Garhett Wyatt, Taylor R Rutherford, Jennifer K Richer, Weston W Porter, Traci R Lyons

## Abstract

Approximately 70% of all breast cancers are estrogen receptor positive (ER+BC) and endocrine therapy has improved survival for patients with ER+BC. Yet, up to half of these tumors recur within 20 years. Recurrent ER+BCs develop resistance to endocrine therapy; thus, novel targets are needed to treat recurrent ER+BC. We found that semaphorin 7A (SEMA7A) confers significantly decreased patient survival rates in ER+BC. We show that *SEMA7A* is hormonally regulated in ER+BC, but its expression does not uniformly decrease with anti-estrogen treatments. Additionally, overexpression of SEMA7A in ER+ cell lines drives increased *in vitro* growth in the presence of estrogen-deprivation, tamoxifen, and fulvestrant. In *in vivo* studies, we found that SEMA7A confers primary tumor resistance to fulvestrant and, importantly, induced lung metastases. Finally, we identify pro-survival signaling as a therapeutic vulnerability of ER+SEMA7A+ tumors and propose that targeting with inhibitors of survival signaling such as venetoclax may have efficacy for treating SEMA7A+ tumors.

**SIGNIFICANCE:** We report that SEMA7A predicts for, and likely contributes to, poor response to standard-of-care therapies and suggest that patients with SEMA7A+ER+ tumors may benefit from alternative therapeutic strategies.

## INTRODUCTION

Breast cancer (BC) is the most commonly diagnosed cancer and the second leading cause of cancer-associated death in women in the United States. Estrogen receptor positive (ER+) BCs comprise approximately 70% of all BC cases. The development of effective anti-estrogen-based endocrine therapies, including aromatase inhibitors (AI) and selective estrogen receptor modulators (e.g., tamoxifen) or degraders (e.g., fulvestrant/Faslodex) has facilitated the successful treatment of ER+ primary tumors. However, depending on nodal involvement, 22-52% of women with ER+BC will experience disease recurrence within 20 years (1), frequently in a distant metastatic site, such as lung, liver, bone, or brain. These recurrent ER+BCs uniformly develop resistance to endocrine therapies, rendering most therapeutic interventions in the clinical setting ineffective at preventing metastatic outgrowth. Therefore, novel molecular targets are needed to predict for and treat recurrent and/or metastatic ER+BCs.

Semaphorin 7a (SEMA7A) is a glycosylphosphatidylinositol-anchored member of the semaphorin family of signaling proteins. SEMA7A has well-described roles in the promotion of axon guidance, immune modulation, and cellular migration (2–6). In adults, SEMA7A is expressed in the brain, bone marrow, lung, and skin, but is minimally expressed in many adult tissues, including the mammary gland (information via https://www.proteinatlas.org). Interestingly, in neuronal cells of the hypothalamus, SEMA7A expression is regulated by steroid hormones (7), while in pulmonary fibrosis models, SEMA7A is regulated by TGF-β (8). In its normal physiological role, SEMA7A binds β1-integrin to activate downstream signaling cascades, including pro-invasive MAPK/ERK and pro-survival PI3K/AKT pathways (2,5,8). Our group and others have described tumor-promotional roles for SEMA7A in mammary cancers, including tumor and host derived mechanisms of increased tumor growth, cellular migration, epithelial to mesenchymal transition, immune cell infiltration, remodeling of the blood and lymphatic vasculature, and metastatic spread (9–13). More recently, it has been reported that SEMA7A expression in lung cancer confers resistance to EGFR tyrosine kinase inhibition (14).

Studies to date have investigated the role of SEMA7A in BC using pre-clinical models of triple negative BC (TNBC), which lack expression of ER (ER-), progesterone receptor (PR-), and amplification of HER2 (HER2-). Specifically, our studies in TNBC reveal that SEMA7A promotes TNBC cell growth, invasion, lymphangiogenesis, lymph vessel invasion, and metastasis (10,15). However, our previous genomics analysis in patient datasets demonstrated that *SEMA7A* expression correlates with poor prognosis in patients with both ER- and ER+BC and that *SEMA7A* expression levels were highest in luminal A tumors, which are generally ER+ and/or PR+ and HER2-(9). Thus, understanding the role of SEMA7A in ER+BC is critical for a complete understanding of its role and impact on BC progression. Our results presented herein show that *SEMA7A* mRNA expression is highest in patients with ER+ BC and that SEMA7A induces pleiotropic tumor-promotional effects in ER+BC cells. We also describe a novel relationship between hormone signaling and expression of SEMA7A in BC. Additionally, we report for the first time that expression of SEMA7A confers resistance to standard-of-care endocrine therapy and to CDK 4/6 inhibition in pre-clinical ER+ models. Finally, using pre-clinical models, we identify a therapeutic vulnerability of SEMA7A-expressing tumors with the BCL2 inhibitor, venetoclax (ABT-199/GDC-0199). These data provide promising evidence that SEMA7A+ER+BCs that do not respond to endocrine therapies may respond to novel treatment regimens in the clinic.

## MATERIALS AND METHODS

### Cell culture

MCF7 cells were obtained from K. Horwitz (University of Colorado Anschutz Medical Campus). T47D and BT474 cells were obtained from V. Borges and M. Borakove (University of Colorado Anschutz Medical Campus). MCF10DCIS cells were obtained from K. Polyak and A. Marusyk (Harvard University, Cambridge, MA). MDA-MB-231 cells were obtained from P. Schedin (Oregon Heath and Sciences University, Portland OR). Tamoxifen resistant T47D cells were obtained from A. Ward and C. Sartorius (University of Colorado Anschutz Medical Campus). Cell lines were validated by fingerprinting analysis at the University of Colorado Anschutz Medical Campus Molecular Biology Core. Cells were routinely tested for mycoplasma contamination throughout the studies described herein with either Lonza MycoAlert PLUS (Walkersville, MD) or Mycoflour Mycoplasma Detection Kit (Molecular Probe, Eugene OR). Cells were cultured at 37 °C and 5% CO2. MCF7 and BT474 cells were grown in MEM with 5% heat-inactivated fetal bovine serum (FBS; Atlanta Biologicals, Flowery Branch, GA) and 1% penicillin/streptomycin (P/S; ThermoFisher Scientific). T47D and E0771 cells were grown in RPMI 1640, with 10% heat-inactivated FBS and 1% P/S. Experiments were performed with cells under passage 30. Forced suspension cultures were performed by coating plates with 12 mg/ml poly-2-hydroxyethyl methacrylate (Sigma, St. Louis, MO). For exogenous SEMA7A treatments, SEMA7A was purified from SEMA7A-Fc expressing MDA-MB-231 cells in collaboration with the CU Anschutz Medical Campus Protein Purification/MoAB/Tissue Culture Core.

### Transfections

For knockdown studies, SEMA7A-targeting shRNA plasmids (shSEMA7A KD1 and KD2) or a scrambled control (Scr; SA Biosciences, Frederik, MD, and the Functional Genomics Facility at the CU Anschutz Medical Campus) were transfected using X-tremeGENE transfection reagent (Millipore Sigma, St. Louis, MO) as described by the manufacturer. SEMA7A overexpression in MCF7 cells was achieved using a SEMA7A-Fc plasmid (generous gift from R. Medzhitov, Yale University, New Haven, CT). A pcDNA3.1 empty vector control plasmid was obtained from H. Ford (CU Anschutz Medical Campus). All other plasmids were obtained from the Functional Genomics Core at the University of Colorado Anschutz Medical Campus. Knockdown and overexpression were confirmed via qPCR and Western blot analysis.

### Luciferase Reporter Assay

The reporter construct was generated by cloning a −820 to +182 base pair fragment of the *SEMA7A* promoter into the pGL3-Basic Luciferase reporter (Invitrogen). The four mutations for the ERE1/2 sites were as follows: TGACCC −>TGTTCC, GGGTCC −> GCATCC, GGGTCA −> GCATCA, GGGCGG −> TTACAG. Three SP1 sites were mutated: GGGCGG −> TTACAG. 50,000 cells were seeded in a 12-well plate and incubated for 5 minutes with 50μL Opti-MEM (Invitrogen, Carlsbad, CA) and 3μL GeneJuice (Millipore Sigma, Burlington, MA) per well. 0.5-1μg of DNA was added to GeneJuice Transfection Reagent/serum-free medium mixture. Cells were harvested 48 hours after transfection using Reporter Lysis Buffer (Promega, Madison, WI), unless treated with 10nM E2. If treated with E2, cells were serum starved for 48 hours in serum-free medium. Cells were then treated with 10nM E2 for 48 hours and harvested as stated above. Luciferase activity was normalized to total protein.

### Drug and Hormone Treatments

Fulvestrant, 4-hydroxytamoxifen, and 17-β estradiol (E2) were purchased from Millipore Sigma (St. Louis, MO). For *in vitro* hormone treatments, cells were cultured in charcoal-stripped serum (Thermo Scientific, Waltham, MA) containing phenol red-free media for 3-5 days, then treated with 10nM E2 as determined by our dose response and the literature(16–18). For clonogenic and growth assays, fulvestrant and 4-hydroxytamoxifen treatments were at a final concentration of 7.5nM and 5μM, respectively. Palbociclib and SU6656 were acquired from MedChemExpress (Monmouth Junction, NJ) and treatments were at a final concentration of 2.5μM and 5μM, respectively, in DMSO. LY294002 was purchased from Cell Signaling Technologies (Danvers, MA) and *in vitro* treatments were at a final concentration of 2.5μM LY294002 in DMSO. *In vitro* venetoclax (Abbvie, Chicago, IL) was at a final concentration of 300nM in DMSO.

### Cell Growth, Invasion, and Death Assays

100,000 MCF7 cells were seeded per well of a 6-well dish to assess growth *in vitro*. Cells were fixed with 10% NBF and stained with crystal violet at the indicated time points. 1,000 MCF7 cells or 5,000 T47D cells were seeded per well of a 6-well dish and assessed 7-14 days later for clonogenic assays. 3D matrigel and collagen cultures were performed as previously described (9,19). Cleaved caspase 7 was measured using a Caspase Glo assay (Promega, Madison, WI) as described by the manufacturer. Exogenous SEMA7A treatments were at a final concentration of 5μg/mL.

### RNA isolation

RNA was isolated using the commercially available Quick-RNA kit (Zymo Research, Irvine, CA). RNA concentration and purity were assessed using a Nanodrop machine via absorbance at 280 nm, 260 nm, and 230 nm. 1 μg of RNA was used to synthesize cDNA with the Bio-Rad (Hercules, CA) iScript cDNA Synthesis kit with the following conditions: 25°C for 5 min., 42°C for 30 min., and 85°C for 5 min.

### mRNA and protein expression analysis

cDNA was quantified via qPCR using Bio-Rad iTaq Universal SYBR Green Supermix and primers for *SEMA7A* (Bio-Rad PrimePCR, Hercules, CA) using the following conditions: 95°C for 3 min, then 40 cycles of 95°C for 15 sec and 60°C for 1 min, 95°C for 30 sec, and 55°C for 1 min. Primer fidelity was assessed for each qPCR experiment using a 95°C melt curve. *SEMA7A* mRNA was detected with Bio-Rad primers (forward: CTCAGCCGTCTGTGTGTATT, reverse: CTCAGGTAGTAGCGAGAGTTTG). Primers for *ESR1 (forward: 5-GAACCGAGATGATGTAGCCA, reverse: GTTTGCTCCTAACTTGCTCTTG*) and *PGR* (*forward: ATTTCATCCACGTGCCTATCC, reverse: CCTTCCTCCTCCTCCTTTATCT*) were purchased from IDP. mRNA quantification was normalized using the geometric average of two reference genes: *GAPDH* (forward: CAAGAGCACAAGAGGAAGAGAG, reverse: CTACATGGCAACTGTGAGGAG) and *RPS18* (forward: GCGAGTACTCAACACCAACA, reverse: GCTAGGACCTGGCTGTATTT). Data were analyzed using the relative standard curve method.

Western blots were performed by loading equal amounts of protein lysates as determined by Bradford assay. Samples were separated on a 10% tris-glycine SDS polyacrylamide gel and electroblotted onto a polyvinylidene fluoride membrane. SEMA7A blots were probed with anti-SEMA7A (clone C6, Santa Cruz Biotechnologies, Dallas, TX) and goat-anti-mouse secondary (Abcam, Cambridge, MA). Blots were then exposed and developed on film. Color images were converted to black and white for presentation, and, if applied, color corrections were applied uniformly to all lanes of the blot to ease viewing of bands. No nonlinear intensity or color adjustments to specific features were performed. Multiple replicates were performed for each blot, and representative blots with corresponding quantifications normalized to loading controls are shown where appropriate.

### Animal models

2,500,000 SEMA7A overexpressing or empty vector control MCF7 cells were injected into the #4 mammary fat pads of 6-7 week old female, estrogen supplemented (20), NSG mice (Jackson Laboratory, Bar Harbor, ME) or NCG mice (Charles River, Wilmington, MA). Tumors were measured using digital calipers every 3-4 days, and tumor volume was calculated as (L x W x W x 0.5). For estrogen independent studies, animals received sham beeswax pellets. For drug treatment studies, fulvestrant dosing began when average tumor volume reached ~200mm^2^ with the following dosing regimen: 2.5mg fulvestrant in 10% ethanol and 90% sesame oil, injected subcutaneously every 5 days. This dose is equivalent to approximately 83mg/kg per mouse and, using the FDA Dosing Guidance for Industry: Conversion of Animal Doses to Human Equivalent Doses, equivalent to 6.7mg/kg in humans. For venetoclax studies, venetoclax was dissolved in phosal-50G:polyethylene glycol-400:ethanol at 6:3:1 to a final concentration of 12mg/mL and administered via oral gavage. Tumors were harvested at designated study endpoints. Mammary glands with intact tumor and lungs were harvested, formalin fixed, and paraffin embedded for immunohistochemical analysis. All animal studies were performed under the approval of the University of Colorado Anschutz Medical Campus Institutional Animal Care and Use Committee.

### Immunohistochemistry

Tissues were formalin-fixed and paraffin-embedded as previously described (21). Antibody information is provided in STable 1. For staining quantification, slides were scanned into the Aperio Image Analysis software (Leica, Buffalo Grove, IL). The total tumor area was quantified by removing necrotic and stromal areas. Custom algorithms for specific stains were used to quantify stain intensity or percent positive stain as assessed by percent weak, medium, and strong determined using the color-deconvolution tool.

### Data Mining Analyses

Kaplan Meir Plotter (22) was used to assess relapse-free survival (RFS) data for 5,143 BC patients with mean follow-up of 69 months. SEMA7A was queried and stratified into high and low expression groups using the auto select best cutoff level function. The generated data were exported, graphed, and analyzed in Graphpad Prism to calculate hazard ratio (HR) with 95% confidence intervals and log rank P values. The Gene Expression Based Outcome for Breast Cancer Online platform (http://co.bmc.lu.se/gobo/) was used to query *SEMA7A* expression in 1,881 available sample tumors and a panel of 51 BC cell lines. Data were stratified as high and low SEMA7A based on median expression and reported by molecular subtype. Outcomes data were reported only in patients with ER+BC. The Oncomine Platform (https://www.oncomine.org) was used to query *SEMA7A* in available BC datasets. The p-value threshold was set to 0.05. All fold-changes and gene ranks were accepted. Data are reported from the molecular subtype analysis, correlation with stage and grade, and drug sensitivity screens.

### IncuCyte Assay

5,000 SEMA7A overexpressing or empty vector control MCF7 cells were seeded per well in a 96 well plate and allowed to adhere overnight. Media was replaced with fresh drug or vehicle treated media the next morning. Drug treatments included 7.5nM fulvestrant, 5μM 4-hydroxytamoxifen, 2.5μM palbociclib, 10μM LY294002, 5μM Su6656, and growth was assessed in an IncuCyte live cell analysis system at the University of Colorado Cancer Center Tissue Culture Core. Confluence was reported every 4 hours.

### Generation of Drug Resistant and Estrogen Deprived Cell Lines

Fulvestrant resistant cell lines were generated by injecting 2,500,000 SEMA7A overexpressing or empty vector control MCF7 cells were injected into contralateral #4 mammary fat pads of a 6 week old female, estrogen supplemented (20) NCG mouse (Charles River, Wilmington, MA). Fulvestrant treatments began 7 days later with the following dosing regimen: 2.5mg fulvestrant in 10% ethanol and 90% sesame oil, injected subcutaneously every 5 days. 48 days after implantation, tumors were harvested and minced in digestion mixture: MCF7 media with 500units/mL collagenase type II and IV, 20μg/mL DNase (Worthington Biochemical, Lakewood, NJ), and 1X Antibiotic-Antimycotic (ThermoFisher, Waltham, MA). Disassociation mixture was incubated at 37°C for one hour. Dissociated cells were centrifuged at 1500 RPM for 7 minutes, then resuspended and plated in standard culture media with 1X Antibiotic-Antimycotic and returned to the incubator overnight. The next day, 7.5nM fulvestrant was added to the media and cells were maintained in these culture conditions. *Ex vivo* fulvestrant resistance was confirmed with clonogenic assays as described above.

Long-term estrogen deprived (LTED) cells were cultured in phenol red-free media (MEM for MCF7 and RPMI-1640 for T47D) supplemented with 5% dextran charcoal stripped FBS for 6 months.

### Drug titration studies

4,000 SEMA7A overexpressing or empty vector control MCF7 cells were seeded per well in a 96 well plate and allowed to adhere for 6 hours. Vehicle or increasing concentrations of 4-hydroxytamoxifen or fulvestrant were added. Growth was assessed in BioSpa live cell imaging system (BioTek, Winooski, VT). Confluence was reported every 4 hours.

### ELISA

Estradiol ELISA kit (Cayman Chemical, Ann Arbor, MI) was used to measure serum estradiol levels from estrogen supplemented or sham beeswax pellet control mice according to the manufacturer instructions.

### Flow Cytometry

MCF7 cells were enzymatically dissociated in Accutase cell detachment solution [Stem cell technology: cat# 07920] for approximately 10 minutes at 37°C. Single cells were stained with anti-mouse/human CD44-AF647 [Biolegend: cat# 103018] and anti-human CD24-FITC [Biolegend: cat# 311104] for 30 minutes at 4°C in the dark. Cells were analyzed on the YETI-ZE5 flow cytometer and data were analyzed using Kaluza software. Cells were delineated from debris by SSC-A/FSC-A and singlets identified by FSC-H/FSC-A. Single cells were subsequently assessed for CD44/CD24 expression.

### Statistical analysis

All cell line and xenograft experiments were performed in biological duplicate or triplicate. Data are expressed as means ± SEM for animal studies and ± SD for *in vitro* studies. All statistical analyses were performed on raw data. An unpaired two-tailed student’s t test was performed for analyses comparing two groups, while a one-way ANOVA followed by a Tukey-Kramer test was used for multiple comparisons of groups of 3 or more. For tumor volume data and incucyte analyses, a repeated measures one-way ANOVA was used. For Kaplan-Meier analyses, the p-value was determined using a log-rank test. All p-values less than or equal to 0.05 were considered statistically significant, and all statistical analyses were performed in GraphPad Prism 7.01 (San Diego, CA).

## RESULTS

### SEMA7A is expressed in ER+ breast cancers and confers poor prognosis

To determine if SEMA7A is expressed in ER+BC, we assessed *SEMA7A* mRNA expression using the Gene Expression-based Outcome for Breast Cancer Online (GOBO) tool. We observed that 67% of luminal subtype cell lines, which are typically ER+, exhibit increased *SEMA7A* expression versus 36-50% of the basal subtypes (which are typically ER-; Figure 1A). Average expression was higher when ER+ cell lines were compared to TNBC cell lines and when luminal cell lines were compared to basal subtype (Figure 1B). *SEMA7A* mRNA expression is also increased in ER+ patient tumors (Figure 1C), and we confirmed that *SEMA7A* was overexpressed and/or had a higher copy number rank in ER+ or PR+ disease in 7 additional datasets using Oncomine (STable 2). Additionally, *SEMA7A* was highest in stage III and IV, grade 2 and 3, and lymph node positive (LN+) disease in multiple datasets (STable 3). We next examined a role for *SEMA7A* status in predicting outcomes for patients with ER+BC; patients with high tumor expression of *SEMA7A* had decreased overall and distant metastasis free survival (Figure 1D-E and SFigure 1A-B). Additionally, using KM Plotter, we further stratified relapse-free survival by LN positivity in the ER+(HER2+/-) subset and observed that high SEMA7A is associated with decreased relapse free survival in patients with both LN+ and LN-status (Figure 1F-G). In fact, the hazard ratio for patients whose tumors were ER+(HER2+/-) SEMA7A+ and with LN-status was the highest. We then confirmed SEMA7A protein expression in ER+ lines (MCF7, T47D, BT474), ER-lines (DCIS, MDA-MB-231, SUM159), and breast epithelial cells derived from human fibrocystic disease (23) (MCF10A) via immunoblot analysis. SEMA7A expression was observed in two of the three ER+ cell lines and one of the three ER-when compared to MCF10A cells (SFigure 1C).

**Figure 1:**
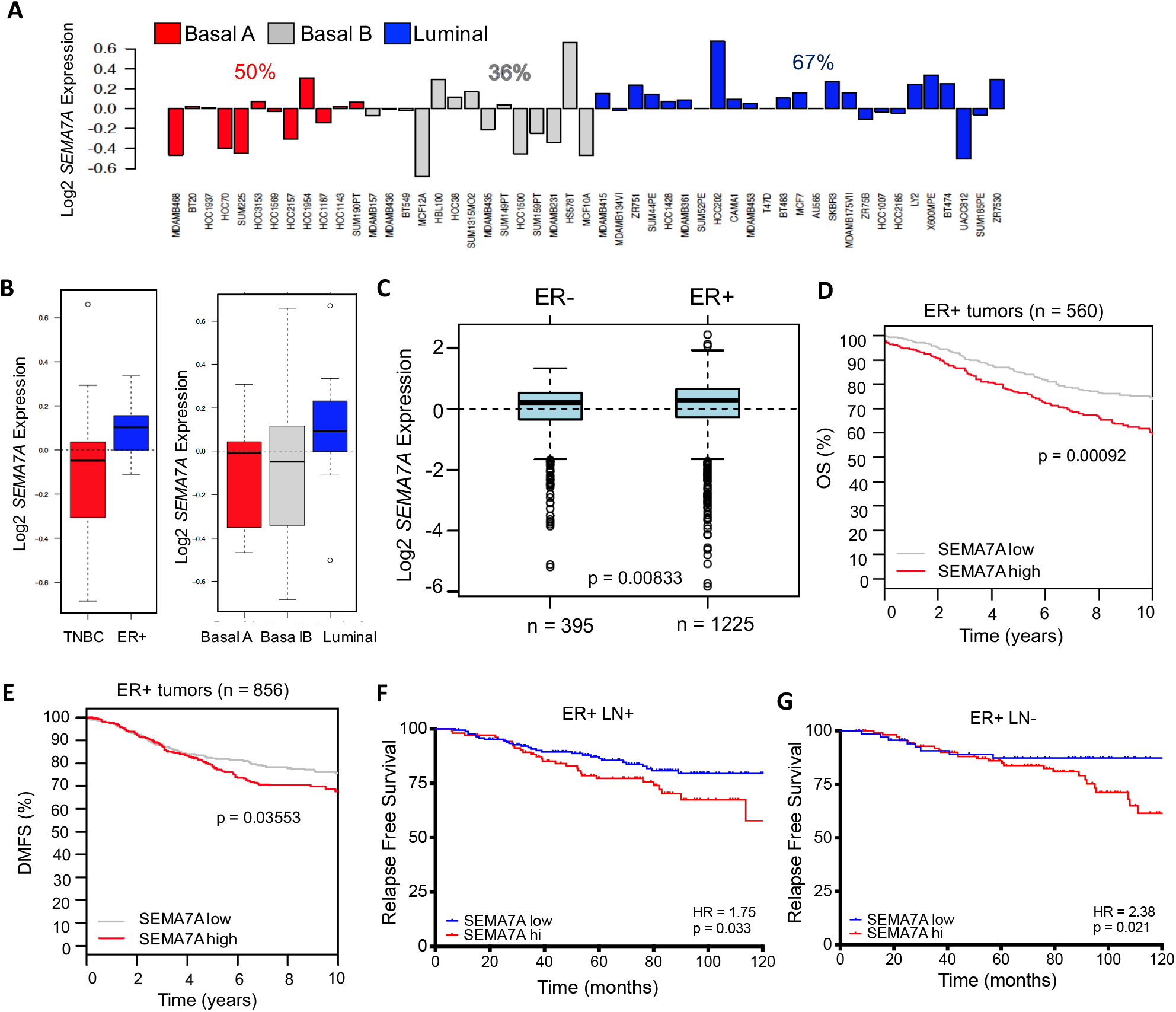
*SEMA7A* is expressed in ER+ breast cancer and associated with poor prognosis. **(A-B)** The Gene Expression-Based Outcome dataset was assessed for *SEMA7A* mRNA expression levels in basal/triple negative breast cancer (TNBC) and luminal/ER+ breast cancer cell lines (n = 51) and **(C)** in ER- or ER+ patient tumors (n = 1620). In the ER+ subset, **(D)** overall survival (OS) and **(E)** distant metastasis free survival (DMFS) curves are stratified by *SEMA7A* high (red) and low (grey) expression. **(F)** Relapse free survival curves for patients with ER+ and PR+ disease and either lymph node positivity (LN+; n = 277) or **(G)** negativity (LN-; n = 185) are stratified by high *SEMA7A* (red) and low *SEMA7A* (blue) using KMplotter mRNA gene chip data. Hazard ratios (HR) are included.

### *SEMA7A* is regulated by estrogen in breast cancer cells and correlates with relapse in patients with ER+ breast cancer

To test the hypothesis that SEMA7A expression could be induced by estrogen (E2) stimulation, we treated ER+BC cells in estrogen deprived conditions with increasing, physiologically relevant (24), concentrations of estrogen. In both MCF7 and T47D cells, E2 resulted in increased SEMA7A protein expression (Figure 2A-B). Then, to determine whether *SEMA7A* expression could be promoted by ER dependent transcription, we performed an *in silico* analysis of the *SEMA7A* promoter using the ConTra (conserved transcription factor binding site) web server (25). We identified two highly conserved estrogen response element (ERE) half sites adjacent to two specificity protein 1 (SP1) consensus binding sites (Figure 2C). SP1 has been shown to facilitate ER binding to ERE half sites in gene promoter regions to promote transcription of estrogen target genes (26); thus, we then utilized a *SEMA7A* luciferase reporter construct to measure promoter activity with E2 treatment and observed increased luminescence with 10nM E2 (Figure 2D). We also observed increased *SEMA7A* mRNA after treatment with 10nM E2 (SFigure 2A). Next, we performed luciferase reporter assays with constructs containing point mutations in the ERE and SP1 sites to show that both are required for induction of *SEMA7A* promoter activity by E2 (Figure 2E-F). To determine whether SEMA7A expression was dependent upon ER, we treated MCF7 and T47D cells with fulvestrant, which decreased expression, but was not sufficient to completely block expression of SEMA7A under normal growth conditions (Figure 2G). Then, to mimic conditions resulting from AI therapy, we tested whether long term estrogen deprivation (LTED) of MCF7 and T47D cells would result in downregulation of SEMA7A. Surprisingly, the cells that were deprived of estrogen exhibited marked upregulation of SEMA7A compared to the wildtype (WT) controls (Figure 2H). Additionally, T47D cells that are tamoxifen resistant (TAMR) exhibit upregulation of SEMA7A (Figure 2I). These results suggest that E2 induces *SEMA7A* expression, but that *SEMA7A* expression can also bypass estrogen dependence. Consistent with this, we observe that MCF7 cells, which normally require estrogen supplementation for growth *in vivo*, can grow in the absence of estrogen supplementation when SEMA7A is overexpressed (Figure 2J; SFigure 2B-C).

**Figure 2:**
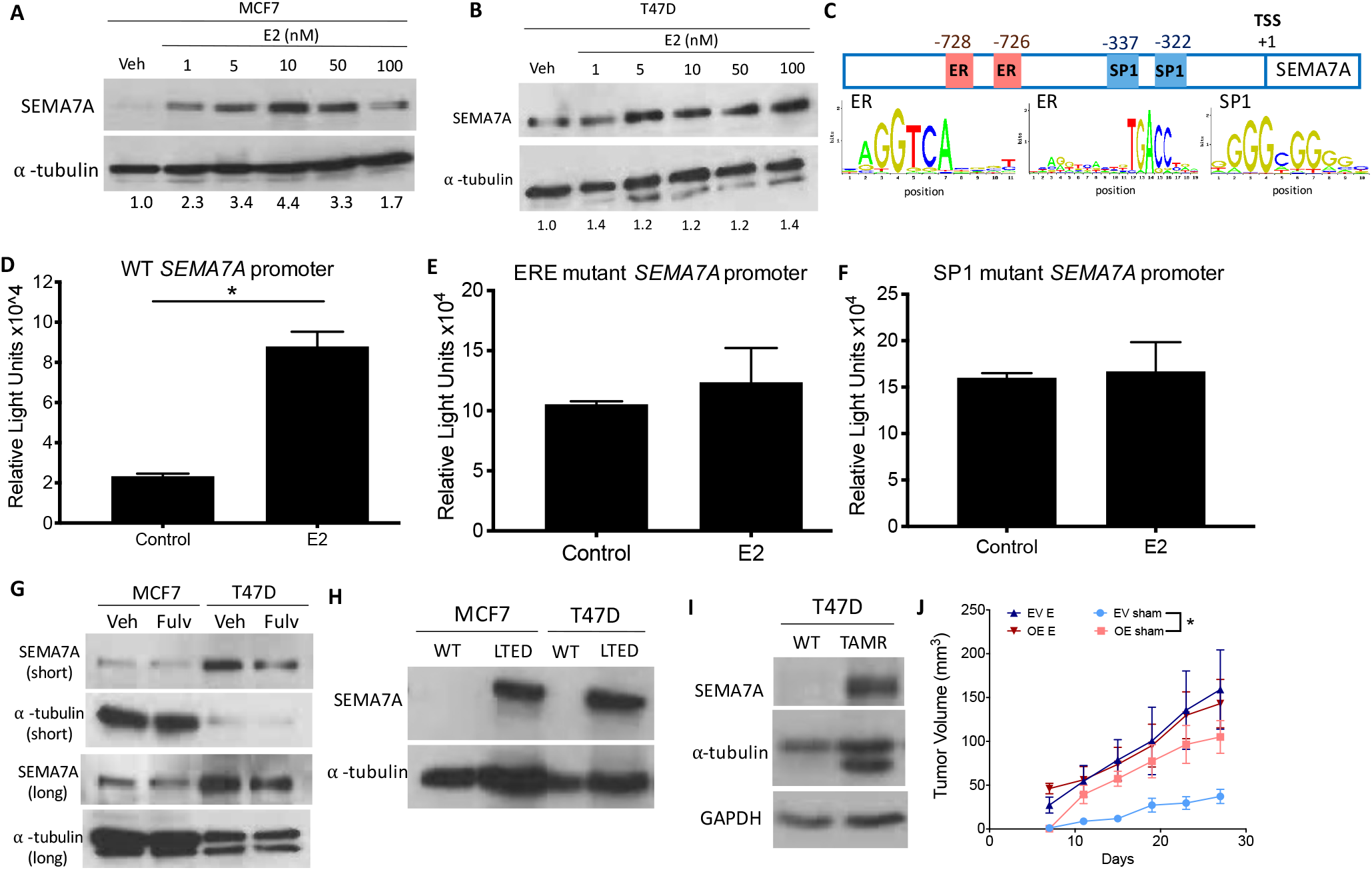
Expression of *SEMA7A* is hormonally regulated. **(A)** MCF7 and **(B)** T47D cells were grown in charcoal stripped conditions for 3 days, then treated with vehicle or increasing concentrations of estrogen (E2). Representative immunoblots for SEMA7A and α-tubulin are shown (n = 3). Normalized quantifications are shown under each blot. **(C)** A ConTra analysis revealed conserved ER and SP1 binding sites in the *SEMA7A* promoter. Distance from the transcriptional start site (TSS) is noted. **(D)** Luciferase activity in MCF7 cells transfected with a luciferase reporter containing the wildtype (WT) *SEMA7A* promoter, **(E)** an estrogen response element (ERE) mutant *SEMA7A* promoter, or **(F)** an SP1 binding site mutant *SEMA7A* promoter upstream of the luciferase gene, then treated with10nM estrogen or vehicle control (n = 3 for each panel). **(G)** MCF7 and T47D cells were treated with 7.5nM fulvestrant for 72 hours. Representative SEMA7A and α-tubulin levels are shown via immunoblot with short and long film exposures (n = 3). **(H)** Representative SEMA7A and α-tubulin levels are shown via immunoblot for WT or long term estrogen deprived (LTED) MCF7 and T47D cells (n = 3), and **(I)** WT or tamoxifen resistant (TAMR) T47D cells (n = 3). **(J)** Tumor volumes for empty vector (EV) or SEMA7A overexpressing (OE) MCF7 tumors in hosts with estrogen pellets (E) or sham controls (n = 6/group). *p < 0.05 by unpaired two-tailed t-test (D) or ANOVA with Tukey’s multiple comparison test (J)

### SEMA7A induces growth and confers therapeutic resistance in ER+ breast cancer cells

To determine if SEMA7A promotes ER+BC progression via increasing tumor cell viability or proliferation, we engineered MCF7 cells to overexpress *SEMA7A*. We confirmed a 3.6-fold increase in SEMA7A protein (SEMA7A OE; Figure 3A) and observed increased cell growth in two-dimensional cultures (Figure 3B), clonogenic assays (Figure 3C), and increased BrdU incorporation in a 3D organoid system that we have previously characterized to model tumor cell growth and invasion into the surrounding tissue (27), which contains Matrigel to model the basement membrane and collagen, which is required for tumor cell invasion (Figure 3D). We also generated two independent shRNA mediated knockdown (KD) variants of SEMA7A and confirmed decreased SEMA7A protein in MCF7 cells (Figure 3E; SFigure 2D). We observed that SEMA7A KD cells were less confluent over a 72 hour time course (Figure 3F), exhibited decreased colony formation in a clonogenic assay (Figure 3G) and SEMA7A KD 3D organoids have significantly reduced proliferation via BrdU incorporation (Figure 3H).

**Figure 3:**
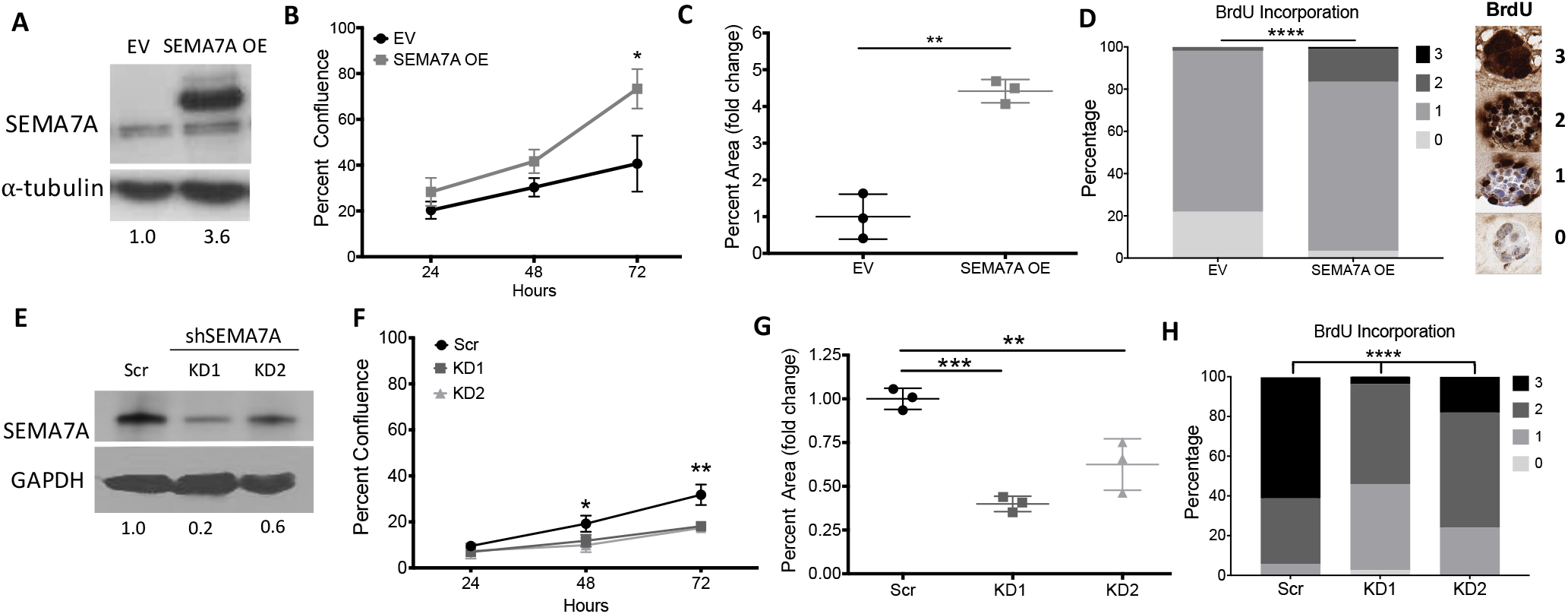
SEMA7A promotes growth of ER+ cells in 2D and 3D cultures. **(A)**. Representative immunoblot for SEMA7A and α-tubulin in MCF7 cell transfected with an empty vector (EV) control or SEMA7A overexpressing (SEMA7A OE) plasmid. Normalized quantifications are shown below (n = 3). **(B)** Confluence of EV or SEMA7A OE MCF7 cells at 24, 48, and 72 hours post seeding (n = 3). **(C)** Clonogenicity of EV or SEMA7A OE MCF7 cells. SEMA7A OE values are normalized to EV controls (n = 3). **(D)** BrdU incorporation was assessed in EV and SEMA7A OE MCF7 cells embedded in Matrigel plus 20% collagen after 5 days in culture. Each organoid received a score 0-3 based on the percent positive of each organoid, where 0 = no BrdU+ cells, 1 = 1-49% BrdU+ cells, 2 = 50-99% BrdU+ cells, and 3 = 100% BrdU+ cells per organoid. Representative BrdU scoring system is depicted on right. **(E)** Representative immunoblot for SEMA7A and GAPDH in MCF7 cell transfected with a with scrambled control (Scr) or shSEMA7A knockdown (KD1 and KD2) constructs. Normalized quantifications are shown below. **(F)**. Confluence of Scr, KD1, or KD2 MCF7 cells at 24, 48, and 72 hours post seeding (n = 3). **(G)** Clonogenicity of Scr, KD1, or KD2 MCF7 cells. KD1 and KD2 values are normalized to Scr controls (n = 3). **(H)** BrdU incorporation was assessed in Scr, KD1, or KD2 MCF7 cells embedded in Matrigel plus 20% collagen after 10 days in culture. Scoring system as described in panel D. *p < 0.05, **p<0.01, ***p < 0.005, ****p < 0.001 by unpaired two-tailed t-test (B-D) or ANOVA with Tukey’s multiple comparison test (F-H)

Kaplan Meier analysis of patients treated with tamoxifen revealed high expression of *SEMA7A* correlated with decreased relapse free survival (Figure 4A). To determine if tamoxifen was ineffective for SEMA7A expressing cells, we cultured MCF7 SEMA7A OE cells in the presence of 4-hydroxytamoxifen. We sought to determine an effective (IC50) dose of 4-hydroxytamoxifenin both EV and OE cells, and we identified a dose, 5μM, that decreased EV cell viability in full serum and in estrogen depleted conditions (SFigure 3A-B). We observed that SEMA7A OE cells grew significantly better than EV controls with 4-hydroxytamoxifen treatment (Figure 4B; SFigure 3C). We also performed a drug titration for fulvestrant and identified 7.5nM as sufficient to decrease MCF7 EV growth (SFigure 3D). We then applied this dose to SEMA7A OE cells. These fulvestrant-treated SEMA7A OE cells grew as well as vehicle-treated EV controls over time and in clonogenic assays (Figure 4C-D; SFigure 4A-B). Additionally, exogenous SEMA7A increased clonogenic outgrowth of both MCF7 and T47D cells, despite the presence of fulvestrant (Figure 4E; SFigure 4C). We further observed that MCF7 SEMA7A OE cells can grow *in vitro* in estrogen depleted conditions (SFigure 4D-E). We next tested how SEMA7A OE tumors grow *in vivo* with endocrine therapy treatments. We chose fulvestrant as we were able to achieve an IC50 dose in the presence of estrogen (SFigure 3D). We implanted MCF7 EV and SEMA7A OE, cells, allowed tumors to establish, then began administering fulvestrant when the tumors reached 200mm^3^. Fulvestrant slowed the growth of control tumors, while the SEMA7A OE tumors continued to grow at the same rate (Figure 4F; SFigure 4F). At study completion, we verified SEMA7A overexpression by immunohistochemistry (IHC; SFigure 5A). Consistent with a drug resistant phenotype, SEMA7A OE tumors exhibited increased proliferation as measured by Ki67 staining at study completion (Figure 4G; SFigure 5B) and were overall less necrotic (Figure 4H; SFigure 5C). Interestingly, ER+ tumor metastases frequently lose expression of ER under conditions of long-term estrogen deprivation (28), and our SEMA7A OE tumors exhibited decreased ER staining intensity by semi-quantitative IHC (Figure 4I-J), which could be a sign of ER independence and explain the lack of response to fulvestrant. Furthermore, *ESR1* and *PGR—* a known ER target gene—downregulation were confirmed in our MCF7 SEMA7A OE cells (SFigure 5D). These results suggest that SEMA7A expression in ER+BC cells renders them less reliant on ER signaling for growth, which also renders them resistant to endocrine therapy and able to disseminate and seed metastases in the presence of tamoxifen, AIs, or fulvestrant. Consistent with this, we observe increased lung metastasis in the SEMA7A OE group (Figure K-L).

**Figure 4:**
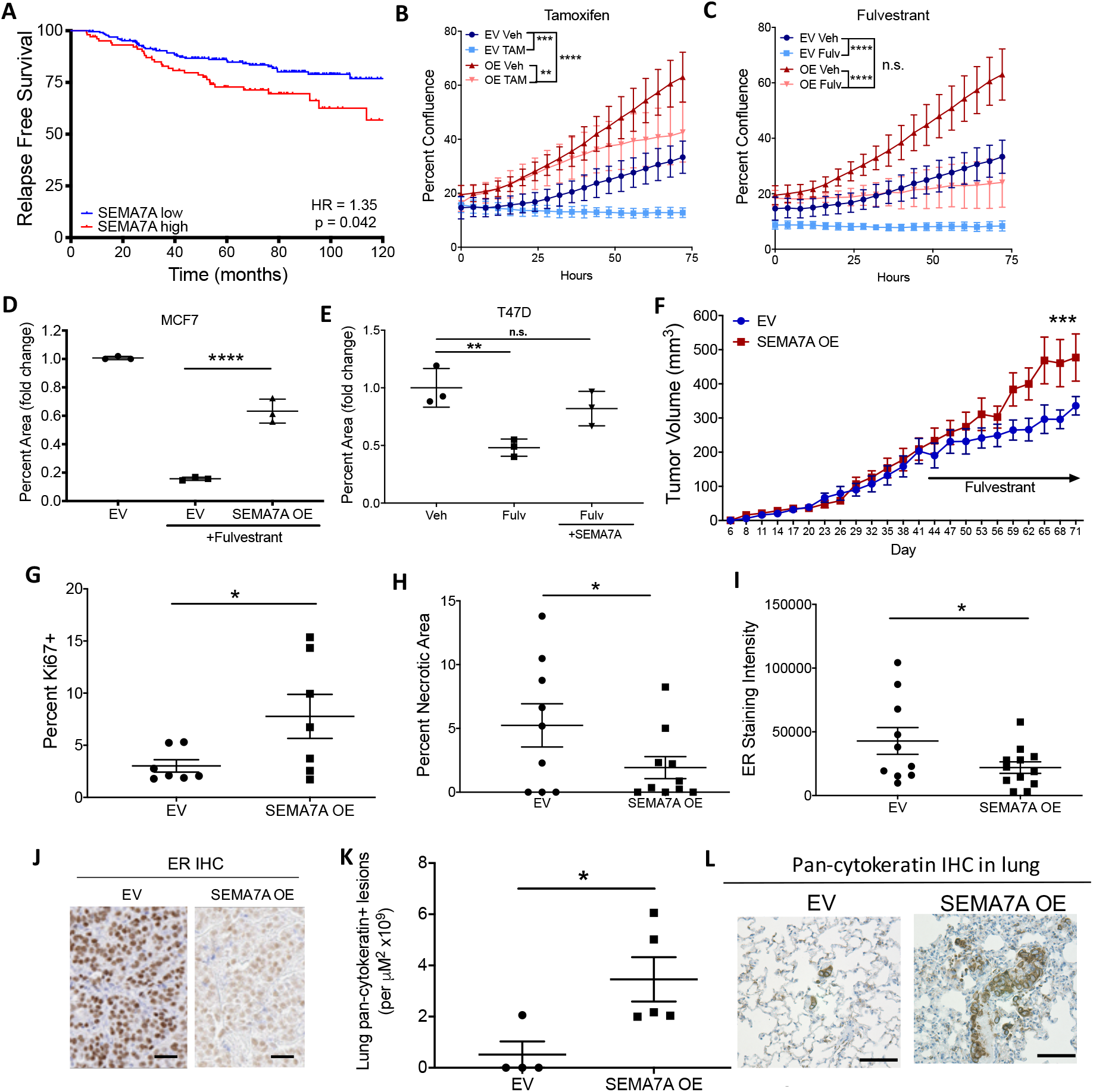
SEMA7A promotes endocrine resistance *in vitro* and *in vivo*. **(A)** Relapse free survival curves for patients with ER+ disease treated with tamoxifen are stratified by high *SEMA7A* (red) and low *SEMA7A* (blue) using KMplotter mRNA gene chip data (n = 335). **(B)** Confluence of MCF7 EV or SEMA7A OE cells treated with 5μM 4-hydroxytamoxifen (n = 3) or **(C)** 7.5nM fulvestrant (n = 3). **(D)** Clonogenic growth of EV or SEMA7A OE MCF7 cells treated with 7.5nM fulvestrant or vehicle. Fulvestrant-treated values are normalized to EV vehicle controls (n = 3). **(E)** Clonogenic growth of T47D cells treated with 7.5nM fulvestrant and 5μg/m L exogenous SEMA7A. Fulvestrant-treated values are normalized to vehicle controls (n = 3). **(F)** Tumor volumes for EV or SEMA7A OE MCF7 tumors in hosts treated with fulvestrant when tumors reached an average of 200mm^3^ (n = 8-10/group). **(G)** Quantification of Ki67, **(H)** necrotic area, and **(I)** ER staining in EV or SEMA7A OE MCF7 tumors in hosts treated with fulvestrant. **(J)** Representative images of ER staining in EV or SEMA7A OE MCF7 tumors. Scale bars 25 μm. **(K)** Quantification and **(L)** representative IHC for pan-cytokeratin in lungs from mice with MCF7 EV or SEMA7A OE tumors (n = 4-6 per group). Scale bars 50 μm. *p < 0.05, **p<0.01, ***p < 0.005, ****p < 0.001 by unpaired t-test (D-I, K) or repeated measures one-way ANOVA with Tukey’s multiple comparison test (B-C)

### SEMA7A induces pro-metastatic phenotypes in ER+ breast cancer cells

Our previous results in ER-BC cells show that SEMA7A induces mesenchymal phenotypes (15), and SEMA7A OE in ER+ MCF7 cells resulted in cells adapting a more mesenchymal morphology (SFigure 5E). We also observed increased invasion of SEMA7A OE MCF7 cells in a transwell filter assay (Figure 5A). We next utilized the 3D model described above where we previously demonstrated that SEMA7A promotes invasion of ER-tumor cells (9,19). Consistent with our hypothesis, SEMA7A OE cells exhibited evidence of invasion as early as 5 days post seeding, when most organoids in the control group remained in rounded, non-invasive spheroid structures (Figure 5B). Conversely, evidence of organoids with invasive protrusions appeared after 10 days in the control cells and were significantly less abundant in SEMA7A knockdowns (Figure 5C).

**Figure 5:**
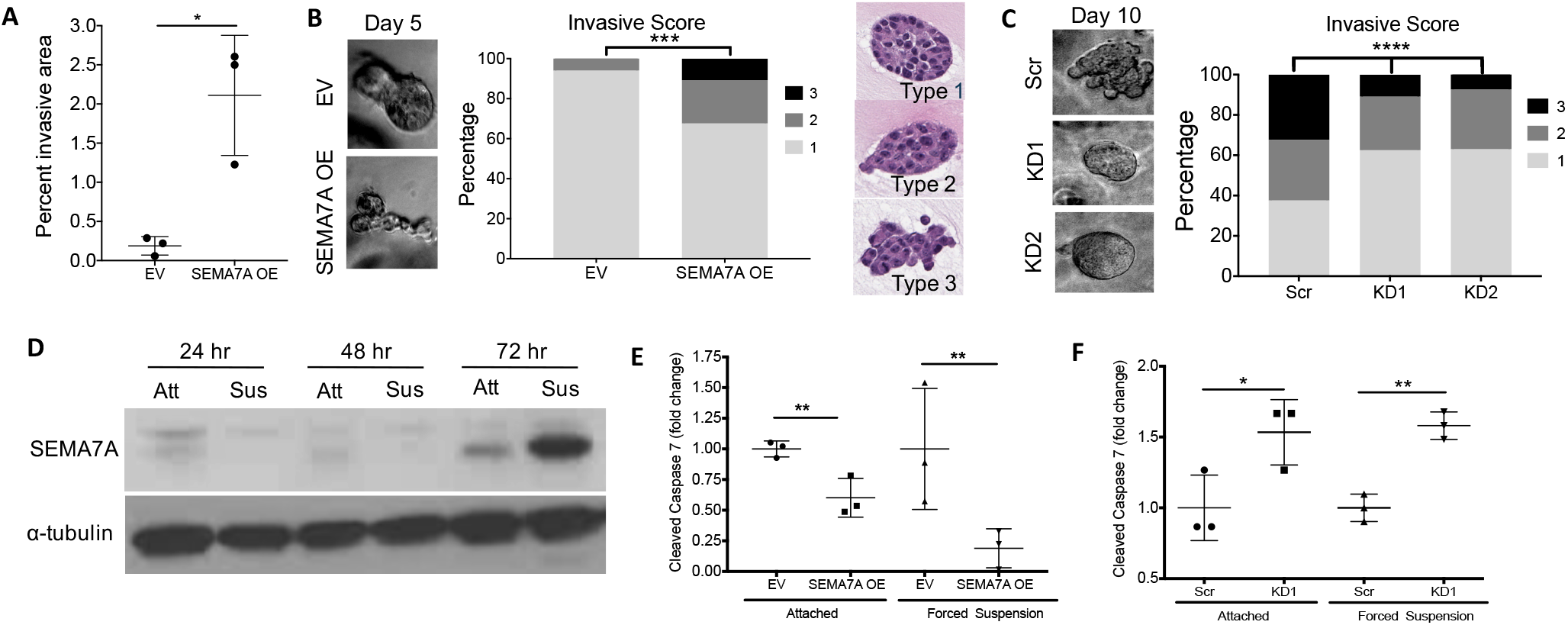
SEMA7A promotes metastatic phenotypes. **(A)**) Percent invasive area is reported from MCF7 EV and SEMA7A OE cells seeded in a transwell invasion assay (n = 3). **(B)** MCF7 EV and SEMA7A OE cells embedded in Matrigel plus 20% collagen scored for invasive phenotypes after 5 days in 3D culture. Type 1 represents non-invasive organoids, type 2 represents moderately invasive organoids, and type 3 represents highly invasive organoids. Representative live images (left), quantification (middle), and H&Es of typing system right) are shown. **(C)** MCF7 Scr, KD1, or KD2 cells embedded in Matrigel plus 20% collagen scored for invasive phenotypes after 10 days in 3D culture. Representative live images (left) and quantification (right) are shown. **(D)** Immunoblot for SEMA7A in wildtype MCF7 cells cultured in normal attached (Att) or forced suspension (Sus) conditions over a timecourse (n = 3). **(E)** Cleaved caspase 7 activity in MCF7 EV and SEMA7A OE (n = 3) or **(F)** Scr and KD1 cells grown in attached or forced suspension cultures for 48 hours (n = 3). Data normalized to EV or Scr controls in attached conditions. *p < 0.05, **p<0.01, ***p < 0.005, ****p < 0.001 by unpaired two-sided t-test (A-B) or one-way ANOVA with Tukey’s multiple comparison test (C, E-F)

After local invasion and intravasation, metastatic tumor cells exit the primary site and survive in non-adherent conditions, which would induce death of normal epithelial cells via a process termed “anoikis” (29). Thus, we utilized forced suspension cultures to determine whether SEMA7A is upregulated in non-adherent conditions. We observed highest SEMA7A expression after 72 hours in suspension, suggesting that SEMA7A may be required for anoikis resistance (Figure 5D). To determine if SEMA7A expression promotes cell survival in forced suspension conditions, we measured cell death via cleaved caspase 7 to reveal decreased cell death in SEMA7A OE cells (Figure 5E). In contrast, KD cells showed a ~2-fold increase in cell death in both attached and forced suspension conditions (Figure 5F), suggesting that SEMA7A may mediate protection from cell death both at the primary site and in circulation during metastasis. Collectively, these data suggest that SEMA7A expression promotes local invasion and cell survival in circulation. This may be one mechanism by which SEMA7A promotes relapse in patients.

### Therapeutic vulnerability of SEMA7A-expressing ER+ tumor cells

Based on our observation that SEMA7A drives growth and metastasis of ER+ tumors and are resistant to endocrine therapy, we postulated that CDK 4/6 inhibitors, which have recently been approved and proven effective for patients with metastatic ER+BC, could inhibit growth of SEMA7A expressing cells. As expected, when treated with 2.5μM of the CDK 4/6 inhibitor palbociclib, MCF7 EV control cells exhibited a significant reduction of growth and clonogenicity *in vitro* compared to vehicle treatment; however, SEMA7A OE cells grew significantly better with palbociclib treatment (Figure 6A-B; SFigure 6A). We therefore sought to identify therapies that may have efficacy for treating patients with SEMA7A-expressing tumors. Using an Oncomine query to identify potential compounds and targetable pathways, we identified 6 compounds predicted to elicit sensitivity of SEMA7A+ tumors, all of which target proteins in either the MAPK or PI3K/AKT signaling pathways (STable 4). Notably, published studies describe these pathways as downstream signaling mediators of SEMA7A activity (2,5,8). Thus, we tested whether growth of SEMA7A OE cells is inhibited by commercially available small molecule inhibitors that target Src and Akt, SU6656 and LY294002, respectively. We observed significantly decreased growth of SEMA7A OE cells with both inhibitors (SFigure 6B-C).

**Figure 6:**
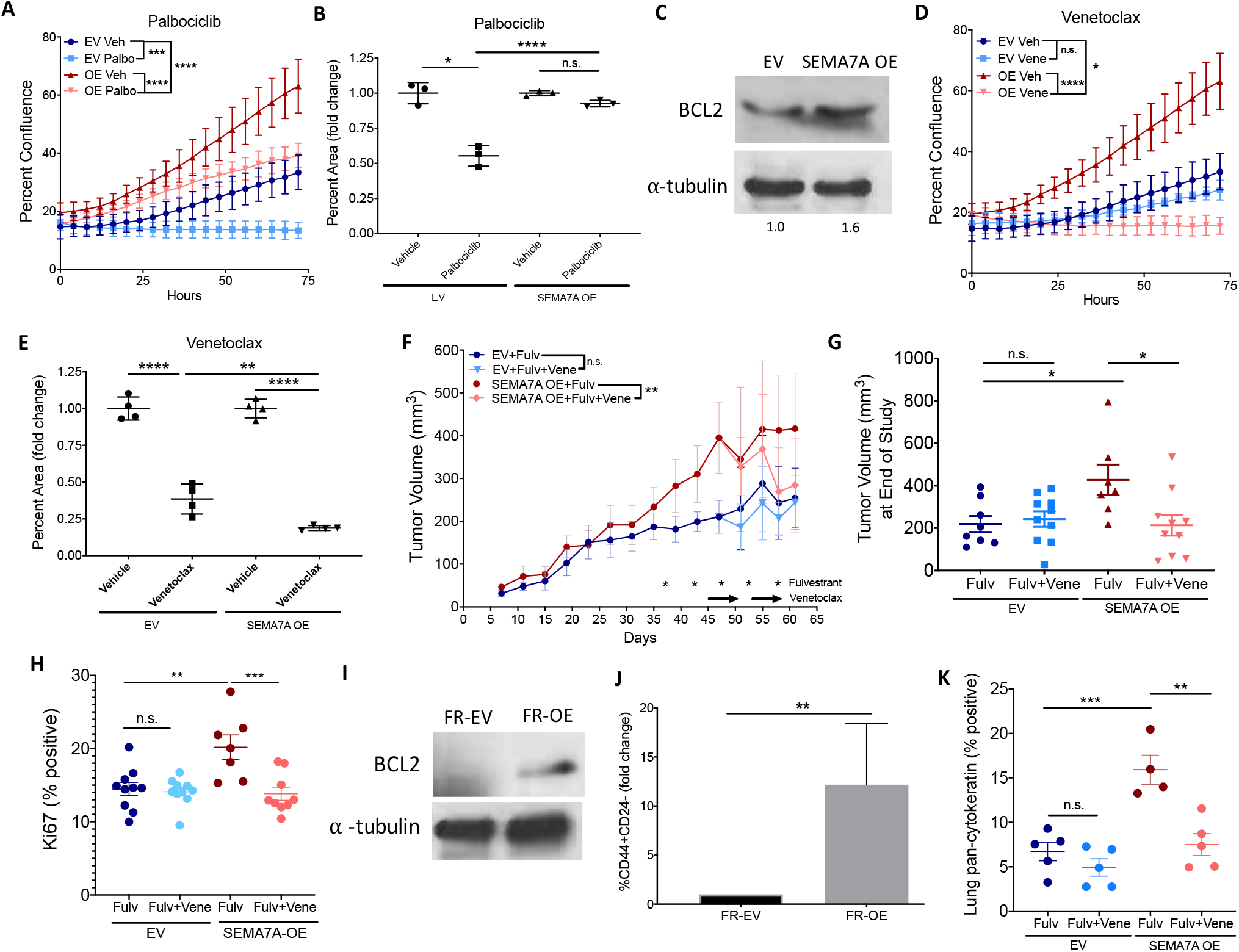
Therapeutic vulnerability of SEMA7A-expressing tumors. **(A)** Confluence and **(B)** clonogenic growth of MCF7 EV or SEMA7A OE cells treated with 2.5μM palbociclib. Clonogenic data are normalized to vehicle controls (n = 3). **(C)** Representative immunoblot of BCL2 and α-tubulin are shown for MCF7 EV and SEMA7A OE cells (n = 3). Normalized quantification is below. **(D)** Confluence and **(E)** clonogenic growth of MCF7 EV or SEMA7A OE cells treated with 300nM venetoclax. Clonogenic data are normalized to vehicle controls (n = 3). **(F)** Tumor volumes of MCF7 EV and SEMA7A OE tumors in hosts treated with fulvestrant +/- venetoclax (n = 4/group). **(G)** Individual tumor volumes of MCF7 EV and SEMA7A OE tumors at end of study after fulvestrant +/- venetoclax treatment. Data are pooled from 2 independent studies. **(H)** Ki67 analysis of MCF7 EV and SEMA7A OE tumors treated with fulvestrant +/- venetoclax. Data are pooled from 2 independent studies. **(I)** Representative immunoblot of BCL2 and α-tubulin are shown for MCF7 fulvestrant resistant (FR) EV and SEMA7A OE cells (n = 3). **(J)** Flow cytometry analysis of MCF7 FR-EV and FR-OE cells. Data are normalized to FR-EV condition. **(K)** Quantification of pan-cytokeratin in lungs from mice with MCF7 EV or SEMA7A OE tumors that were treated with fulvestrant +/- venetoclax. Data pooled from 2 independent studies. *p < 0.05, **p<0.01, ***p < 0.005, ****p < 0.001 by one-way ANOVA with Tukey’s multiple comparison test (B,E-H,K) or repeated measures one-way ANOVA with Tukey’s multiple comparison test (A,D)

However, as these compounds are not clinically available, we sought to identify a compound that would inhibit the downstream pro-survival function induced by these pathways. Venetoclax is an orally available inhibitor of the anti-apoptotic BCL2 protein that has recently demonstrated synergy in combination therapy approaches in patients with ER+BC (30). We first identified that our cells expressed BCL2 and observed increased expression in SEMA7A OE cells (Figure 6C). Then, we tested the sensitivity of SEMA7A OE cells to venetoclax at a clinically relevant dose. Proliferation of SEMA7A OE cells was decreased with venetoclax treatment as compared to vehicle treatment and to EV control cells treated with venetoclax, while clonogenic outgrowth was decreased in both EV and SEMA7A OE cells (Figure 6D-E and SFigure 6D). To test whether fulvestrant resistant SEMA7A OE tumors respond to venetoclax treatment *in* vivo, we implanted MCF7 EV control or SEMA7A OE cells into NCG mice. Tumors were allowed to grow to an average of 200mm^3^, when we then initiated fulvestrant treatment. As expected, the SEMA7A OE tumors continued to grow, while tumor growth plateaued in the control group. After separation of the curves was observed, we administered vehicle or venetoclax treatment in combination with fulvestrant. We observed a dramatic decrease in tumor growth, tumor growth rate, final tumor volume, and decreased Ki67+ tumor cells in the SEMA7A OE, which was surprisingly not observed in the EV control group (Figure 6F-H and SFigure 6E&F). Next, we sought to examine whether BCL2 expression was affected by the fulvestrant treatment, as BCL2 is an ER target gene. To this end, we harvested cells from fulvestrant treated tumors and cultured them *ex vivo* in the presence of fulvestrant (SFigure 7A-B). Immunoblot analysis of these cells revealed that the fulvestrant resistant EV (FR-EV) cells lost expression of BCL2, while the fulvestrant resistant SEMA7A OE (FR-OE) cells maintained expression (Figure 6I). Additionally, the FR-OE cells were enriched for CD44+CD24-cell populations (Figure 6J; SFigure 8A-C), which exhibit increased capacity for self-renewal, generation of heterogenous progeny, enhanced invasive phenotypes and drug resistance (31,32); these results suggest that SEMA7A OE may result in enrichment for drug resistant stem cells in response to endocrine therapy, which may also promote metastasis. To determine whether there is potential for venetoclax to prevent metastasis of SEMA7A+ER+ tumors, we measured lung metastasis in our co-treated animals and found a significant decrease with the addition of venetoclax to fulvestrant treatment (Figure 6K). Thus, our results suggest that venetoclax may be an viable treatment option for inhibiting growth, recurrence, and metastasis of SEMA7A-expressing ER+BCs, which are likely to be resistant to endocrine therapies.

## DISCUSSION

Determining whether patients with ER+BC will or will not respond to endocrine therapy and identifying novel treatment options for patients with endocrine resistant ER+BC is of the utmost importance, as patients with ER+ disease comprise the largest percentage of BC patients and the largest number of BC-related deaths. The semaphorin family of proteins have multiple, and variable, described roles in cancer, with some demonstrating tumor-suppressive functions and others demonstrating tumor-promotional roles (33). The data presented in this study contribute to an emerging body of evidence describing a role for SEMA7A in tumor progression. Our lab and others have recently begun to characterize SEMA7A as a mediator of breast/mammary, melanoma, and lung cancer (9–14). In this report, we describe a novel role for SEMA7A in ER+BC recurrence and metastasis, where SEMA7A expression confers resistance to anti-endocrine therapy and predicts for poor response in patients with ER+ disease. In previously published data, we showed that SEMA7A promotes numerous tumor-promotional phenotypes in ER-preclinical models, including tumor cell growth, motility, invasion, and cell survival (9,15). We also reported that SEMA7A induces macrophage-mediated remodeling of lymphatic vasculature, as well as tumor cell dissemination via the lymphatics, and others have demonstrated SEMA7A-dependent angiogenesis in ER-BC models (10,11).

Motility and invasion are known hallmarks of metastatic cells (29). However, as cells venture through the metastatic cascade, they must also adapt to many different microenvironments such as certain hostile conditions, including anchorage independent conditions during travel through the vasculature or lymphatics. We now show that SEMA7A expression promotes anoikis resistance and consequent anchorage-independent survival and we extend our observations to show that SEMA7A overexpression can promote spontaneous and experimental lung metastasis. Additionally, we suggest that the pleiotropic effects promoted by SEMA7A may give SEMA7A-expressing tumors a selective advantage under the stress of endocrine therapies, which results in long-term hormonal depravation intended to halt tumor cell growth and induce cell death. Our results support that SEMA7A expression allows tumor cells to bypass reliance on ER, rendering them reliant instead upon SEMA7A mediated cell survival. This is further evidenced by our observations that SEMA7A-expressing tumors downregulate expression of ER—a mechanism that warrants further investigation. Furthermore, published studies report that SEMA4C promotes MCF7 tumor growth, viability, metastasis, and renders ER+ cells non-responsive to tamoxifen via downregulation of ER (34), suggesting that this mechanism may extend to other multiple semaphorin family members.

Metastatic disease in patients with ER+BC frequently exhibits endocrine resistance, and many metastases downregulate ER or express mutant forms of the *ESR1* gene. Although the prevalence of ER loss has been debated, a recent meta-analysis reveals approximately 23% of ER+ tumors have corresponding ER-metastases (28). Further, tumors with low expression of ER (1-10% positive) are reported to behave more like ER-tumors, and patients with ER low tumors have significantly lower disease free survival rates compared to patients with ER high (11-100% positive) tumors (35). These results are consistent with our observations that SEMA7A expressing cells exhibit pro-metastatic phenotypes and are also resistance to therapy and downregulate ER. We suggest that initial expression of SEMA7A in hormone receptor positive disease is promoted cooperation between ER and SP1 at the SEMA7A promoter, as we demonstrate that SEMA7A expression can be promoted by estrogen treatment in a dose-dependent manner and that both the ERE half sites and SP1 sites are required for induction of promoter activity by estrogen. However, SEMA7A expression is not completely inhibited with ER degradation in ER+BC cells, suggesting additional mechanisms are at play to regulate SEMA7A expression. Studies of other semaphorins have described hormone receptor regulation. In ovarian cancer, pro-proliferative *SEMA4D* is positively regulated by ER (36). Conversely, expression of *SEMA3B* and *SEMA3F*, which are both growth-suppressive, in ovarian cancer is positively regulated by E2 and progesterone (37). Finally, *SEMA3C* is transcriptionally regulated by the androgen receptor (AR) in prostate cancer cells (38) and promotes growth, migration, and invasion in BC (39), is detectable in metastatic lung cancer cells (40), and promotes survival of gastric cancer and glioma stem cells (41,42). SEMA3C also has a described role in promoting drug resistance in ovarian cancer (43). Consistent with a mechanism for SEMA7A in drug resistance, our observation that ER levels in SEMA7A OE tumors are decreased after fulvestrant treatment suggests that fulvestrant had the intended effect, but that fulvestrant alone was not sufficient to control growth in SEMA7A expressing tumors, suggesting a bypass mechanism. Taken together, these studies support that future investigations, utilizing chromatin immunoprecipitation, will be necessary to show direct promoter binding of SEMA7A by multiple nuclear hormone receptors, including ER, PR and AR.

In addition to endocrine therapy resistance, we demonstrate that SEMA7A expression confers resistance to the CDK 4/6 inhibitor palbociclib, which is used in combination with endocrine therapy for first line treatment of metastatic disease or in subsequent lines following disease progression (44). Our data suggest that patients with SEMA7A+ tumors may be at high risk for relapse and poor response to current standard-of-care therapies, and that novel therapeutic strategies could be applied for patients pre-selected by SEMA7A status. Interestingly, Kinehara et al. recently described that SEMA7A promotes resistance to standard-of-care EGFR tyrosine kinase inhibitors in lung cancer through anti-apoptotic β1-integrin-mediated ERK signaling; SEMA7A was thus proposed as a predictive marker for response to TKI therapy and potential therapeutic target for resistant tumors (14). This report, along with our findings, suggests that SEMA7A may confer resistance to multiple targeted therapies in numerous tumor types; therefore, direct targeting and therapeutic vulnerabilities of SEMA7A+ tumors warrant additional investigation in a clinical setting. Moving toward this goal, results presented herein suggest that SEMA7A+ tumors may be targetable via other mechanisms. Specifically, SEMA7A has demonstrated the ability to activate PI3K/AKT and MAPK/ERK signaling (2,5,8). We now report that SEMA7A OE cells have increased expression of the anti-apoptotic protein BCL2. We therefore tested if SEMA7A+ tumor cells exhibit sensitivity to selective inhibitors of these pathways, including the BCL2 inhibitor venetoclax. Our results showing that SEMA7A+ tumors cells exhibit reduced growth and viability after exposure to these therapeutics support that targeting downstream effectors of SEMA7A may prove an effective strategy for recurrent SEMA7A+ tumors, which are resistant to currently available therapeutic regimes. We also show, for the first time, that when treated with fulvestrant, SEMA7A expressing tumors maintain expression of BCL2 while controls lose BCL2 expression. Previous reports have described increased BCL2 expression in fulvestrant-resistant MCF7 cells, and fulvestrant sensitivity can be restored with a BCL2 inhibitor or BCL2 knockdown (45). Additionally, the first clinical study of the BCL2 inhibitor venetoclax in solid tumors revealed that patients with metastatic ER+BC received great benefit from this drug in combination with tamoxifen, and a recent report describes a benefit of adding venetoclax to fulvestrant/palbociclib in preclinical ER+ models (46). While we have not explored this triple combination in our current studies, future studies will explore whether venetoclax can reverse the resistance to palbociclib that we observe in our SEMA7A OE cells. Since venetoclax is clinically available, administered orally, and well-tolerated, we propose that novel clinical trials could test whether patients with ER+SEMA7A+ cancer would benefit from combination treatment in both the adjuvant and metastatic setting. Finally, clinical trials assessing SEMA4C as a biomarker for BC diagnosis and relapse are already underway (ClinicalTrials.gov identifiers: NCT03663153 and NCT03662633). Since similar phenotypes are observed in pre-clinical models with SEMA4C and SEMA7A, we propose that SEMA7A may also warrant investigation in the clinic as a predictive marker of ER+BC recurrence.

## ACKNOWLEDGEMENTS

The authors acknowledge the CU Anschutz Cancer Center Protein Production/MoAB/Tissue Culture Shared Resource, the Biorepository Core Facility, and the University of Colorado Cancer Center Support Grant (P30CA046934) for SEMA7A protein production, Incucyte use, and Aperio Software use. Contents are the authors’ sole responsibility and do not necessarily represent official NIH views. The authors also thank the following persons for technical support and feedback in the production of this manuscript: Alan Elder, Alexander Stoller, Veronica Wessells, Michelle Borakove, Virginia Borges, Chloe Young, Sarah Tarullo, Annie Pham, Paris Franz, Craig Jordan, and Brett Stevens.

## Author Contributions

LSC and TRL conceived and designed the study. LSC performed all *in vivo* studies. LSC, GLW, and TRR performed all *in vitro* studies. JKR and WWP assisted with methodology development and data interpretation. GLW, WWP, and JKR provided critical reagents. LSC and TRL were responsible for hypothesis development, conceptual design, and data analysis. LSC and TRL wrote the manuscript, and all authors provided critical evaluation, editing, and review of the manuscript.

